# BCG-induced T cells shape *Mycobacterium tuberculosis* infection before reducing the bacterial burden

**DOI:** 10.1101/590554

**Authors:** Jared L. Delahaye, Benjamin H. Gern, Sara B. Cohen, Courtney R. Plumlee, Shahin Shafiani, Michael Y. Gerner, Kevin B. Urdahl

## Abstract

Growing evidence suggests the outcome of Mycobacterium tuberculosis (Mtb) infection is established rapidly after exposure, but how the current tuberculosis vaccine, BCG, impacts early immunity is poorly understood. Here we found that murine BCG immunization promotes a dramatic shift in infected cell types. While alveolar macrophages (AM) are the major infected cell for the first two weeks in unimmunized animals, BCG promotes the accelerated recruitment and infection of lung infiltrating phagocytes. Interestingly, this shift is dependent on CD4 T cells, yet does not require intrinsic recognition of antigen presented by infected AM. Mtb-specific T cells are first activated in lung regions devoid of infected cells, and these events precede vaccine-induced reduction of the bacterial burden, which occurs only after the co-localization of T cells and infected cells. Understanding how BCG alters early immune responses to Mtb provides new avenues to improve upon the immunity it confers.

## Introduction

Bacillus Calmette-Guerin (BCG), the current tuberculosis (TB) vaccine, is effective at preventing disseminated disease in infants and young children (1). However, in most settings it provides little or no protection against adult pulmonary TB, the transmissible form of disease (2). Thus, despite widespread BCG immunization for nearly a century, *Mycobacterium tuberculosis* (Mtb) kills over 1.5 million people every year, more than any other single infectious agent (3). A better TB vaccine is urgently needed, but attaining this goal has been surprisingly difficult (4). Furthermore, because BCG reduces childhood mortality, a new vaccine will likely be added to a regimen that includes BCG, rather than replace it (5). To develop a strategy that builds upon BCG-mediated protection, we must first understand how BCG shapes immunity to Mtb, especially during early stages of infection when protective immunity is established.

In mice, pulmonary Mtb burdens are equivalent between BCG-immunized and control mice until two weeks after infection (6). The failure of BCG to impact the Mtb burden during the first two weeks of infection has been attributed to the delayed arrival of T cells in the lung (7). However, BCG-specific T cells have been shown to be present in the lungs (8) of immunized mice even prior to Mtb challenge, indicating that impaired T cell recruitment cannot fully account for the inability of BCG to induce early protection.

In this study, we utilized the mouse model to investigate the impact of BCG on the early immune response to Mtb infection. Our findings reveal unexpected roles for CD4 T cells in: 1) accelerating the translocation of Mtb-infected alveolar macrophages (AM) into the lung interstitium; 2) recruiting monocyte-derived macrophages; and 3) promoting the early transfer of Mtb from AM to other phagocytes.

## Materials and Methods

### Mice

C57BL/6 and MHCII^−/−^ mice were purchased from Jackson Laboratories (Bar Harbor, ME). All mice were housed in specific pathogen-free conditions at Seattle Children’s Research Institute (SCRI). Experiments were performed in compliance with the SCRI Animal Care and Use Committee. Both male and female mice between the ages of 8-12 weeks were used.

### BCG immunization

BCG-Pasteur was cultured in Middlebrook 7H9 broth at 37°C to an OD of 0.2-0.5. Bacteria was diluted in PBS and 10^6^ CFU in 200ul was injected subcutaneously. After immunization, mice were rested for 8 weeks prior to Mtb infection.

### Aerosol Infections

Infections were performed with wildtype H37Rv Mtb or H37Rv transformed with an mCherry reporter plasmid (9). Mice were enclosed in a Glas-Col aerosol infection chamber and 50-100 CFU were deposited directly into the lungs.

### Intratracheal and intravenous labeling

For intratracheal labeling, 30min prior to sacrifice, mice were anesthetized with 25% isoflurane in propylene glycol (Fisher Scientific) and 0.25ug of CD45.2 PE-Cy7 in 50ul of PBS was pipetted into the airway. For intravenous (i.v.) labeling, mice were anesthetized as above and infused with CD45.2 PE 10 min prior to sacrifice.

### Lung cell isolation and antibody staining

Mouse lungs were homogenized in HEPES buffer with Liberase Blendzyme 3 (70ug/ml; Roche) and DNaseI (30ug/ml; Sigma-Aldrich) using a gentleMacs dissociator (Miltenyi Biotec). Lungs were incubated at 37°C for 30 min and then further homogenized with the gentleMacs. Cells were filtered through a 70um cell strainer and resuspended in RBC lysis buffer (Thermo) prior to a PBS wash. Cells were next incubated with 50ul Zombie Aqua viability dye (BioLegend) for 10min at room temperature. Viability dye was quenched with 100ul of antibody cocktail in 50% FACS buffer (PBS containing 2.5% FBS and 0.1% NaN3)/50% Fc block buffer. Staining was performed for 20min at 4°C. Cells were washed with FACS buffer and fixed with 2% paraformaldehyde for 1hr prior to analysis on an LSRII flow cytometer (BD Biosciences). When stain sets contained tetramers, staining was performed for 1hr at room temperature. Ag85B and TB10.4 tetramers were obtained from the NIH Tetramer Core Facility.

### Imaging

Mice were infected with H37Rv Mtb-mCherry and sacrificed at D10 and D14. Lungs were excised and submerged in BD Cytofix fixative solution diluted 1:3 with PBS for 24hr at 4°C. Lungs were washed 2x in PBS and dehydrated in 30% sucrose for 24hr prior to OCT embedding and rapid freezing in a methylbutane-dry ice slurry. 20um sections were stained overnight at room temperature and coverslipped with Fluoromount G mounting media (Southern Biotec). Images were acquired on a Leica SP8X confocal microscope, compensated for fluorophore spillover using LAS X (Leica), and analyzed with Imaris (Bitplane) and FlowJo (10).

### T cell depletion

Mice were intraperitoneally injected with 400ug anti-CD4 GK1.5 or anti-CD8 2.43 (BioXcell) in PBS at D-1, D4, and D10 relative to infection.

### Bone marrow chimeras

WT CD45.1/2 F1 mice were irradiated with 1000 rads and reconstituted with a 1:1 mixture of CD3-depleted (Miltenyi Biotec) CD45.1 B6.SJL:CD45.2 MHCII^−/−^ bone marrow. At D56 post-reconstitution, mice were immunized with BCG.

### Th1 polarization and adoptive transfers

CD4 T cells from ESAT-6-specific (C7) (11) CD90.1^+^ and OVA-specific (OTII) CD45.1^+^ TCR transgenic mice were negatively enriched from spleens using EasySep magnetic microbeads (STEMCELL). T cells were Th1 polarized as follows: 1.6 × 10^6^ transgenic T cells were cultured with 8.3 × 10^6^ irradiated CD3^−^ splenocytes. 5 μg/ml of ESAT-6 or OVA peptide, 10 ng/ml IL-12, and 10 μg/ml of anti–IL-4 antibody (R&D Systems) were added at D0. At D3, cells were split 1:2, and 10 ng/ml IL-12 was added (R&D Systems). On D5, Th1 cells were i.v. injected into B6 CD45.2^+^ mice infected with Mtb 35 days prior.

## Results and Discussion

### BCG vaccination promotes Mtb egress from AM early in infection

To understand the effects of BCG immunization on early Mtb infection, we examined the pulmonary Mtb burdens in BCG-immunized and control mice. Consistent with prior reports (6, 7), lung burdens rose similarly in both groups through two weeks (Fig. 1A). At D15, the Mtb burden in the immunized group began to diverge and was reduced by one log by D21. These findings are consistent with the idea that BCG-induced immunity is not initiated until the third week of Mtb infection.

**Figure 1:**
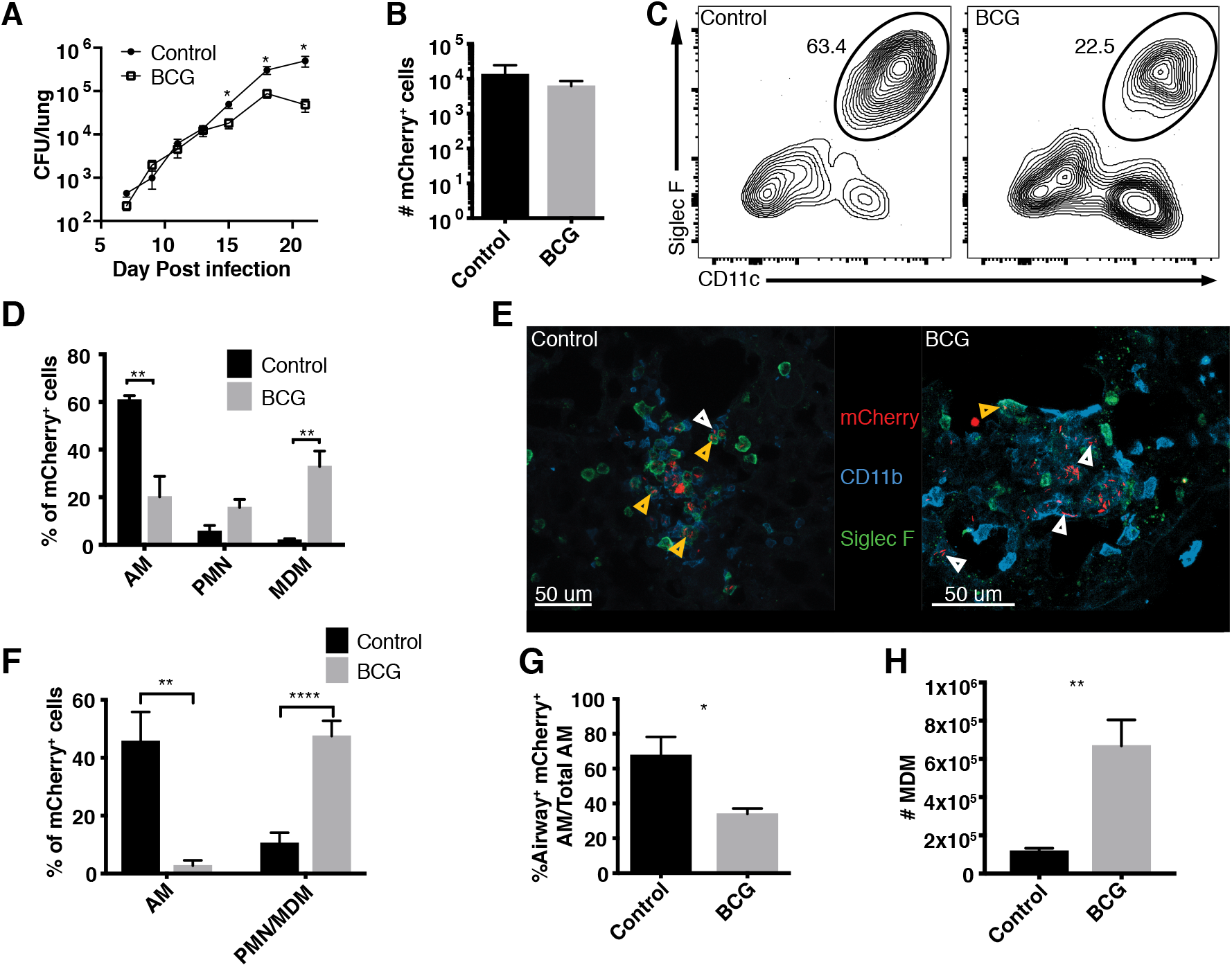
BCG vaccination promotes Mtb egress from AM early in infection. (A) Mtb burden in the lungs of mice that did or did not receive BCG (n=4 mice/group/timepoint). (B) Total number of mCherry^+^ lung cells at D14 by flow cytometry (n=4-5 mice/group). (C) Representative flow plot of the proportion of mCherry^+^ cells identified as CD11c^+^ Siglec-F^+^ AMs at D14. (D) Composition of mCherry^+^ lung cells (AM: CD11c^+^ Siglec-F^+^, PMN: CD11b^+^ Ly6G^+^, MDM: CD11b^+^ CD64^+^) at D14 by flow cytometry (n=4-5 mice/group). (E) Representative images of the lung at D14 showing infected Siglec F^+^ AM (orange arrows) and infected Siglec F^−^ CD11b^+^ cells (white arrows). (F) Composition of mCherry^+^ lung cells at D14 by quantitative histocytometry (n=6-8 infectious foci from 2 mice/group). (G) Ratio of airway label positive infected AM at D14 (n=5 mice/group). (H) Number of MDM in the lung at D14 by flow cytometry (n=4-5 mice/group). Single-group comparisons were performed by unpaired t test. Data are presented as mean ± SEM. *p < 0.05, **p < 0.01, ****p < 0.0001. All experiments were performed at least 2–3 times.

We recently found that Mtb first infects AM before disseminating to other cells including neutrophils (PMN) and monocyte-derived macrophages (MDM) (12). As tissue-resident and recruited phagocytes have been shown to differ in their capacity to curb Mtb replication (13, 14), we next asked whether immunization alters the proportions of cell types that harbor infection. Consistent with the similar Mtb burdens at D14, the numbers of cells harboring fluorescent Mtb (Mtb-mCherry) were also similar in each group (Fig. 1B). Surprisingly, even at this early phase, we observed a dramatic shift in the composition of infected cells. At D14, the proportion of Mtb-infected AM was significantly reduced in immunized animals compared to controls, with a corresponding increase in infected PMN and MDM (Fig. 1C-D). We confirmed these findings using confocal microscopy and quantitative histocytometry (10), wherein most Mtb was within SiglecF^+^ AM at D14 in controls but within CD11b^+^ SiglecF^−^ cells (primarily PMN and MDM) in immunized mice (Fig. 1E-F). As Mtb dissemination to PMN and MDM requires translocation of infected AM to the lung interstitium (12), we next assessed whether this translocation was accelerated in immunized mice. Indeed, intratracheal antibody administration, which specifically labels alveolar-localized cells (12), revealed significantly increased interstitial localization (label-negative) of infected AM in immunized mice at D14 (Fig. 1G). Finally, immunization significantly enhanced MDM recruitment to the lung at D14 (Fig. 1H), which was not observed at earlier time points or in Mtb-naïve mice (Supplemental Fig. 1A), suggesting that the accelerated recruitment of MDM in immunized mice begins between D10 and D14. Thus, although BCG does not impact the pulmonary Mtb burden in the first 2 weeks of infection, it accelerates the translocation of infected AM from alveoli to the lung interstitium, MDM recruitment, and Mtb dissemination to PMN and MDM.

### BCG accelerates the recruitment of antigen-specific T cells to the lung following Mtb infection

This unexpected impact of BCG on the early dynamics of infection led us to next investigate how immunization affects the kinetics of T cell recruitment to the lung. Before infection, antigen-specific CD4 (Ag85B) and CD8 (TB10.4) T cells could be identified in lung cell suspensions of immunized mice (Fig. 2A-C). Although ∼25% of the Ag85B-specific cells were located in the lung parenchyma (as evidenced by their failure to stain with i.v. CD45 antibody), virtually all of the TB10.4-specific cells resided in the vasculature (Fig. 2D). Following infection, immunized mice had significantly more Ag85B-specific and TB10.4-specific cells in the lung parenchyma than controls as early as D10; by D14 they contained >5-fold more (Fig. 2B-C). Thus, BCG induces a small population of lung-resident Mtb-specific CD4 T cells prior to infection. After infection, BCG accelerates the pulmonary recruitment of both CD4 and CD8 Mtb-specific T cells, even before impacting the Mtb burden.

**Figure 2:**
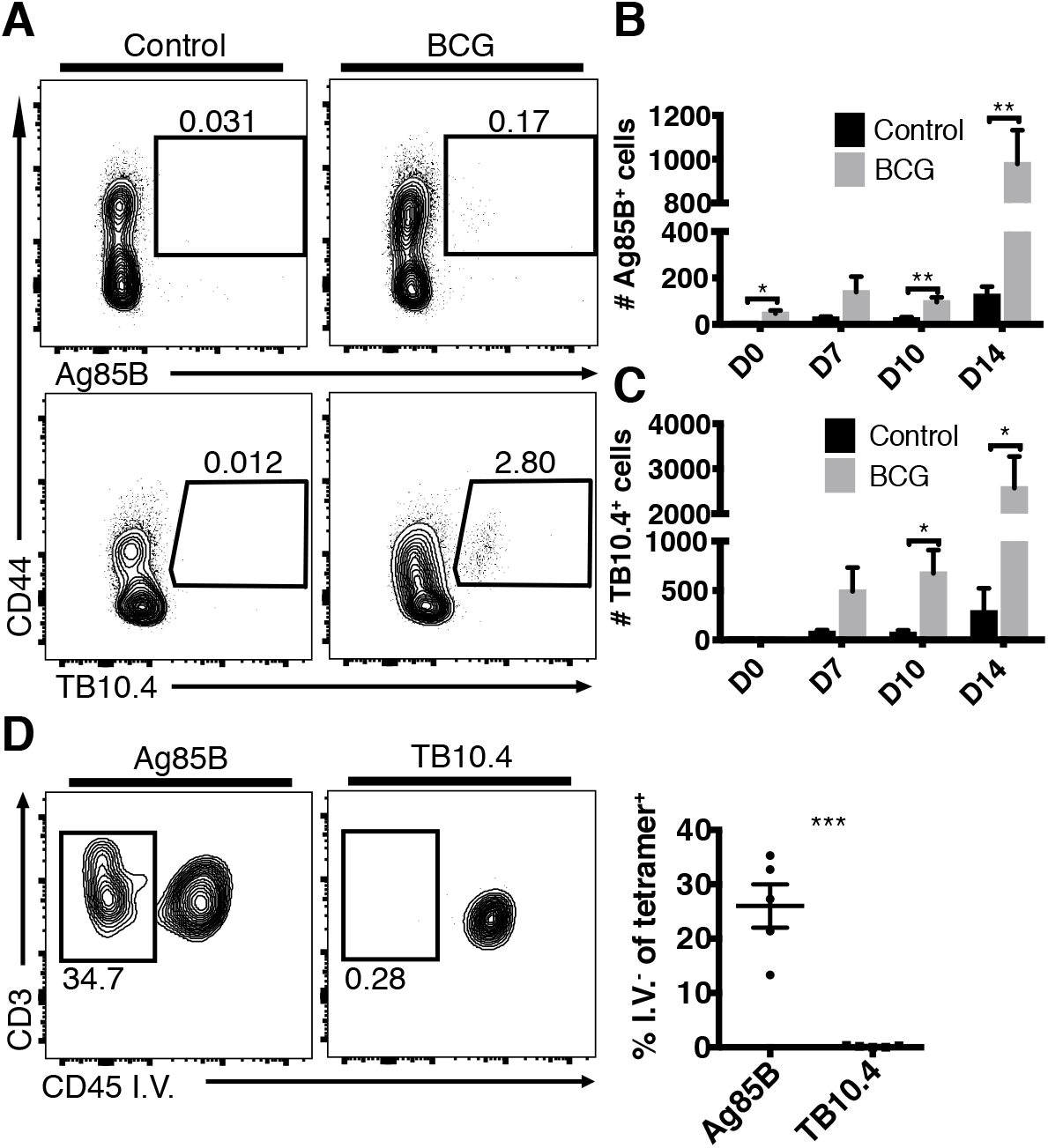
BCG accelerates the recruitment of antigen-specific T cells to the lung following Mtb infection. Time course of the number of tetramer-specific T cells in the lung. Mice received i.v. CD45 antibody prior to sacrifice. (A) Representative flow plots showing Ag85B-specific (CD3^+^CD4^+^) and TB10.4-specific (CD3^+^CD8^+^) T cells in the lungs of control and immunized mice prior to infection. The tetramer^+^ cells in immunized mice are further gated on CD45 i.v.^−^ to determine the proportion in the lung parenchyma. Total number of i.v.^−^ Ag85B-specific (B) and TB10.4-specific (C) cells in the lungs of control and immunized mice (n=3-5 mice/group/timepoint). (D) Proportion of tetramer^+^ cells that are i.v.^−^ in immunized mice at D0 (n=5 mice/group). Single-group comparisons were performed by unpaired t test. Data are presented as mean ± SEM. *p < 0.05, **p < 0.01, ***p < 0.001. All experiments were performed at least twice.

### CD4 T cells are required for the accelerated transfer of Mtb from AM to recruited phagocytes

Given the presence of lung-resident Mtb-specific T cells in immunized mice prior to infection, we next determined whether T cells play a role in the accelerated transfer of Mtb from AM to other myeloid cells. CD4 or CD8 T cells were depleted from immunized mice beginning 1 day prior to Mtb-mCherry infection and lung cells were assessed at D14 (Supplemental Fig. 1B-C). In the absence of CD4 T cells, the accelerated transfer of Mtb from AM to PMN and MDM was partially reversed, whereas CD8 T cell depletion had no effect (Fig. 3A). Interestingly, the accelerated MDM recruitment (Fig. 1H) was also abolished by CD4 depletion (Fig. 3B). We next investigated whether direct recognition of Mtb-infected cells by CD4 T cells was required for the early dissemination out of the AM niche and whether MHCII^−/−^ AM, which cannot present antigen to CD4 T cells, would retain Mtb longer than WT AM. WT:MHCII^−/−^ mixed bone marrow chimeras were generated, BCG immunized, and infected with Mtb-mCherry. At D14, BCG induced the accelerated transfer of Mtb from AM to other myeloid cells irrespective of intrinsic MHCII expression (Fig. 3C). Taken together, BCG-induced CD4 T cells promote the early transfer of Mtb from AM to other myeloid cells in a process that does not require direct cognate interactions between T cells and Mtb-infected AM. Our finding that CD4 T cells promote MDM recruitment to the lung, thereby providing new bacterial targets, may help explain the increased proportion of infected MDM in immunized animals. This recruitment likely relates to T cell production of cytokines, such as IFN-gamma and TNF, which are known to trigger the release of chemokines that act on MDM, i.e., CCL2 and CXCL10 (15).

**Figure 3:**
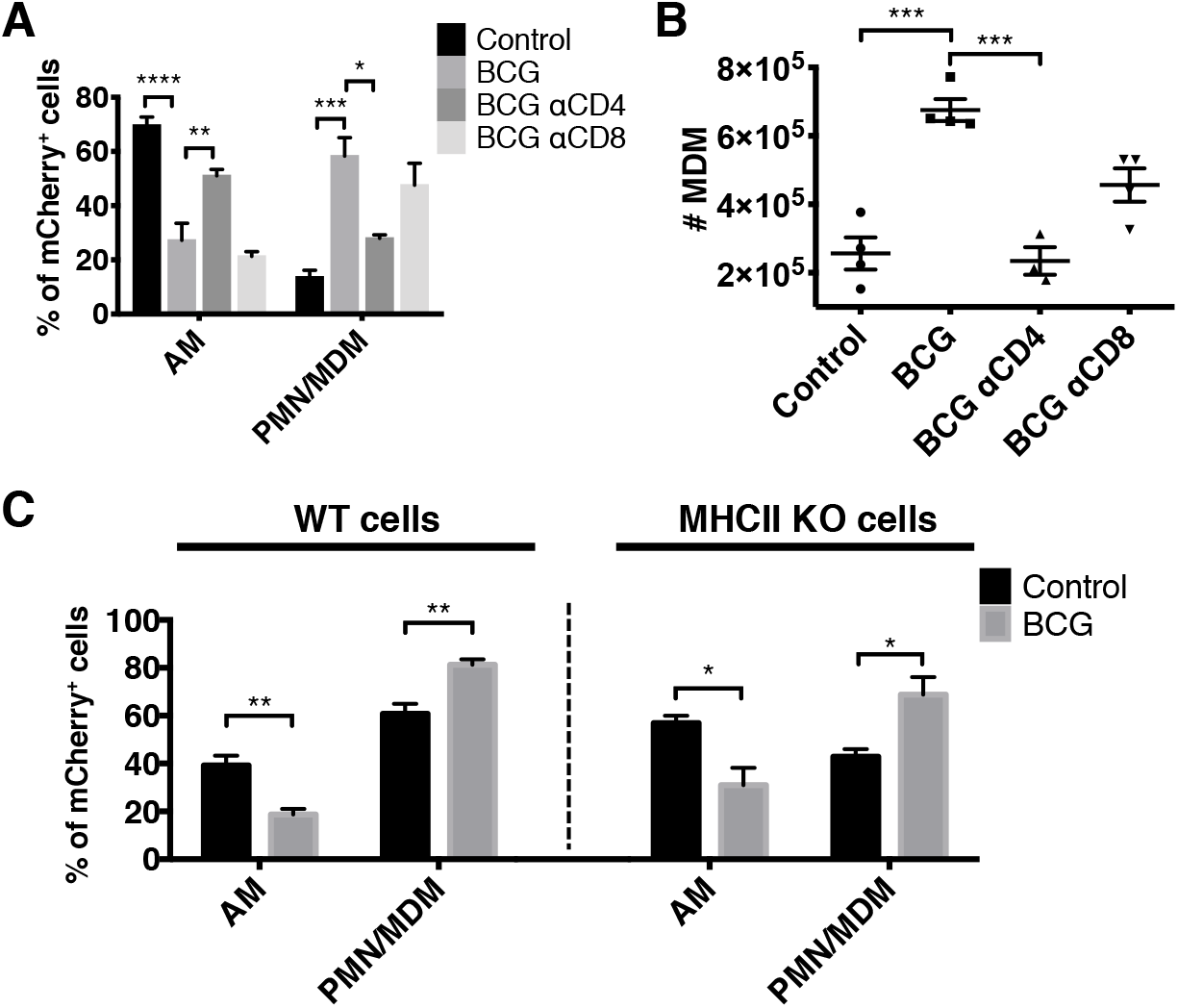
CD4 T cells are required for the accelerated transfer of Mtb from AM to recruited phagocytes. (A) Composition of mCherry^+^ cells in control, immunized, and T cell-depleted immunized mice at D14 (n=4 mice/group). (B) Total number of MDM as in (A). (C) Composition of WT (left) and KO (right) mCherry^+^ cells in control and immunized mixed bone marrow chimeras at D14 (n=3-4 mice/group). Single-group comparisons were performed by unpaired t test (C) and multiple-group comparisons by one-way ANOVA (A and B). Data are presented as mean ± SEM. *p < 0.05, **p < 0.01, ***p < 0.001, ****p < 0.0001. All experiments were performed at least twice.

### BCG-induced CD4 T cells are initially activated distal to the site of Mtb infection

We next investigated the site of CD4 T cell activation during early Mtb infection using phospho-S6 (pS6) as a marker of TCR signaling, which is rapidly induced by TCR engagement, peaking at 4 h and resolving within 24 h (16). To confirm that pS6 expression by T cells is TCR-dependent in the context of Mtb-infected lungs, we demonstrated that pS6 was robustly expressed by adoptively transferred TCR transgenic Mtb-specific (ESAT-6; C7) CD4 T cells compared to irrelevant TCR transgenic T cells (OVA-specific) (Supplemental Fig. 2A). We next performed quantitative histocytometry to assess the intrapulmonary location of pS6 expression by CD4 T cells. At D10, there were significantly more pS6^+^ CD4 T cells in the lungs of immunized mice compared to controls (Fig. 4A-B, Supplemental Fig. 2F). Surprisingly, few of these cells were located near infected cells (Fig. 4A, 4D, Supplemental Fig. 2B-E). Thus, although BCG induces early T cell recruitment and activation, at D10 this occurs primarily in uninfected areas of the lung, which may be due to Mtb antigenic export from infected to uninfected antigen-presenting cells (17). Taken together, the activation of BCG-induced CD4 T cells, which occurs distal to sites of infection, shapes immunity to Mtb challenge earlier than previously appreciated by facilitating the pulmonary recruitment of MDM and accelerating the transfer of Mtb from AM to other myeloid cells. This transfer likely influences the ability of the BCG-immunized host to control Mtb, as prior studies have shown that tissue-resident vs. recruited macrophages differ profoundly in their capacity to control Mtb replication (13, 14). Future studies are needed to elucidate the overall impact on protection because the settings in which distinct macrophage types mediate enhanced immunity remain unclear.

**Figure 4:**
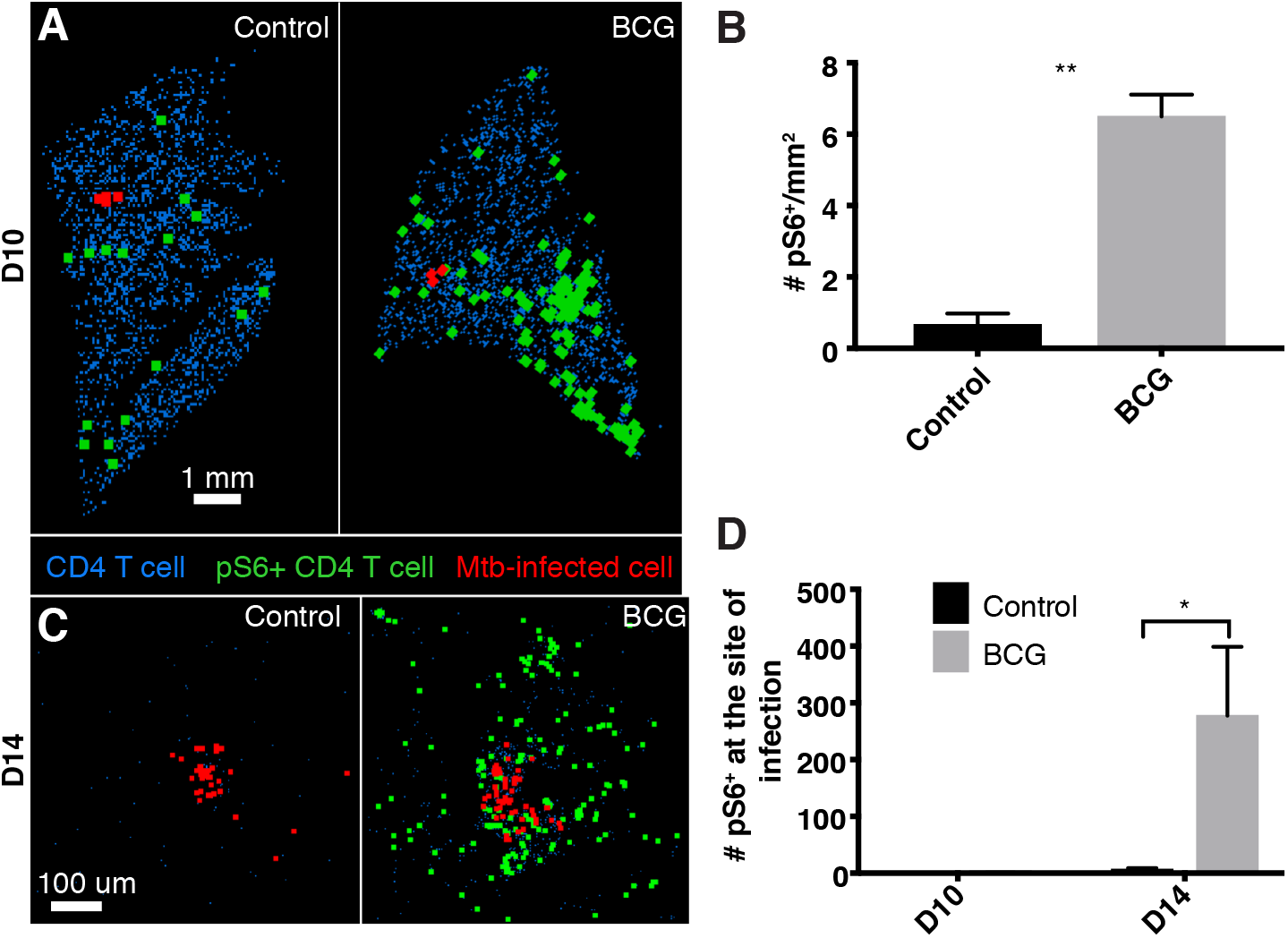
BCG-induced CD4 T cells are initially activated distal to the site of Mtb infection. Quantitative histocytometry was used to identify the location of CD4 T cells (blue) and pS6^+^ CD4 T cells (green) relative to infected cells (red) in lung sections at D10 (A) and sites of infection at D14 (C). (B) Number of pS6^+^ CD4 T cells per mm^2^ of lung at D10 as determined by quantitative histocytometry (n=2-3 mice/group). (D) Number of pS6^+^ CD4 T cells within 80 μm of an infected cell (n=2 mice/group). This cutoff was based on the limit of IFNg diffusion within tissue (21). Single-group comparisons were performed by unpaired t test. Data are presented as mean ± SEM. *p < 0.05, **p < 0.01.

Interestingly, BCG-induced CD4 T cells only begin to curb Mtb replication at D14, when they finally co-localize with cells harboring Mtb, as evidenced by the identification of many pS6^+^ T cells at sites of infection compared to controls (Fig. 4C-D). This is consistent with the finding that optimal immunity against Mtb requires direct interactions between antigen-specific CD4 T cells and Mtb-infected cells (21). Why do T cells and Mtb-infected cells not co-localize earlier? The AM is the first cell type to become infected and remains the primary infected cell type for at least a week (12). During this time, the immune system appears largely unaware of the looming threat, as few MDM or PMN are recruited to the lung. The recent finding that AM infection is non-inflammatory and poorly induces chemokines may help explain the covert nature of early infection (Rothchild, A.C. et al. 2019. bioRxiv: 520791). Furthermore, the replication and spread within the AM population, a process associated with cell death of infected macrophages and phagocytosis by other macrophages, likely involves apoptosis, as necrotic cell death is associated with chemokine release and recruitment of MDM/PMN (19). Perhaps vaccine-induced T cells that express receptors for apoptotic “find-me” signals could co-localize with infected AM and exhibit earlier Mtb control compared to BCG-induced T cells, which may not express such receptors (20). Together, these results further our understanding of the features of pulmonary Mtb dissemination in the context of BCG, which could aid rational vaccine design to effectively complement BCG.

## Acknowledgements

We thank P. Andersen, J. Woodworth, and R. Mortensen for critical feedback of the manuscript as well as the flow cytometry core and vivarium staff for technical assistance.

## Notes

1 This study was supported by the NIH grants 1R01AI134246 (K.B.U.), 1R01AI076327 (K.B.U.), U19AI135976 (K.B.U.), 1K22AI108628-01A1 (M.Y.G.), and T32GM007270-42 (J.L.D.).

