## Supplemental Figures for "BCG-induced T cells shape *Mycobacterium tuberculosis* infection before reducing the bacterial burden"

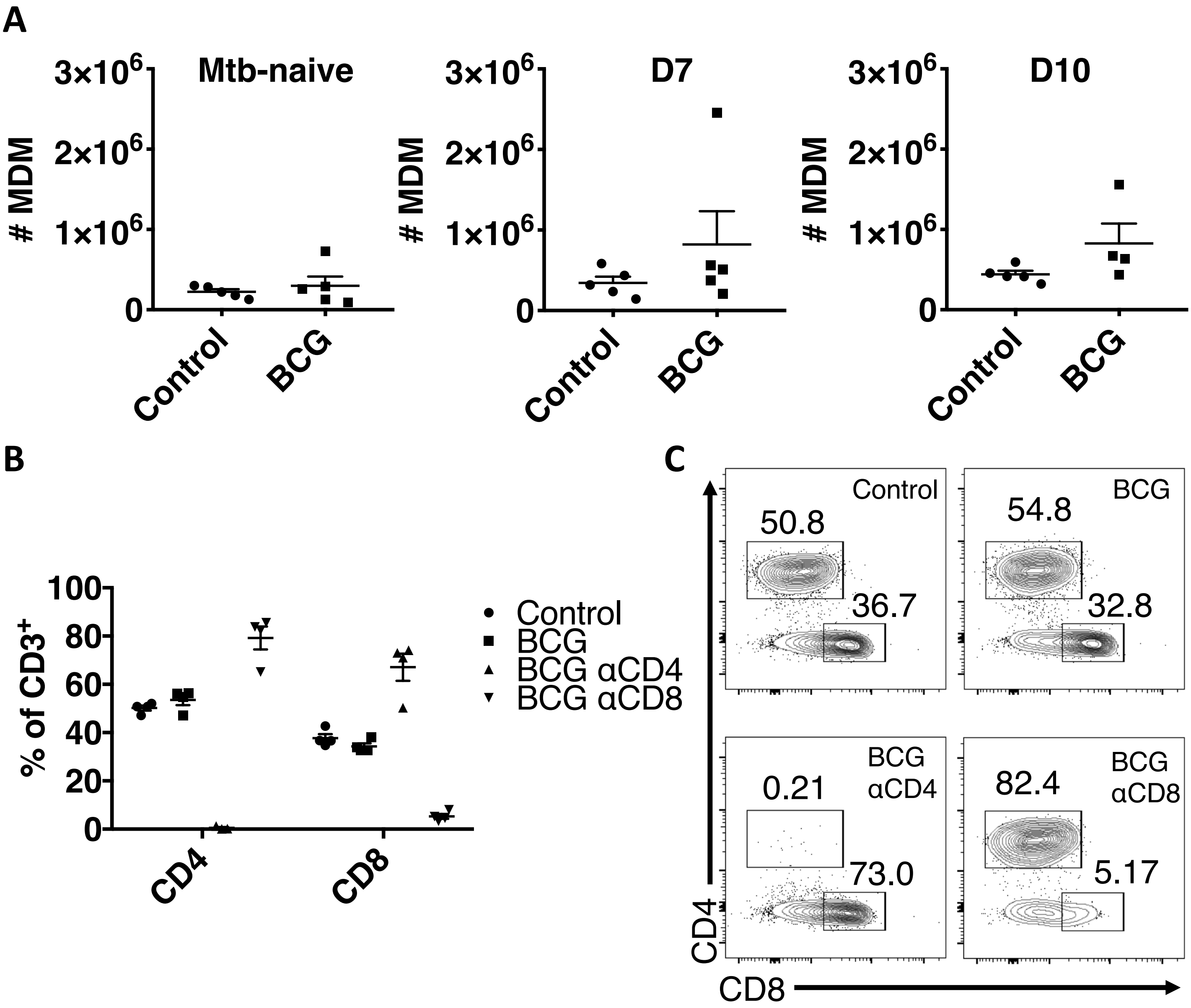


Supplemental Figure 1: Kinetics of MDM recruitment and T cell depletion efficacy.
(A) Quantification of the number of MDM in the lungs of animals prior to and following Mtb infection as determined by flow cytometry (n=4-5 mice/group). Single-group comparisons were performed by unpaired t test. Data are presented as mean ± SEM. (B) Proportion of CD4 and CD8 T cells in control, BCG, and BCG animals that were treated with CD4 or CD8-depleting antibodies (n=4 mice/group). (C) Representative flow plots showing CD4 and CD8 T cells.

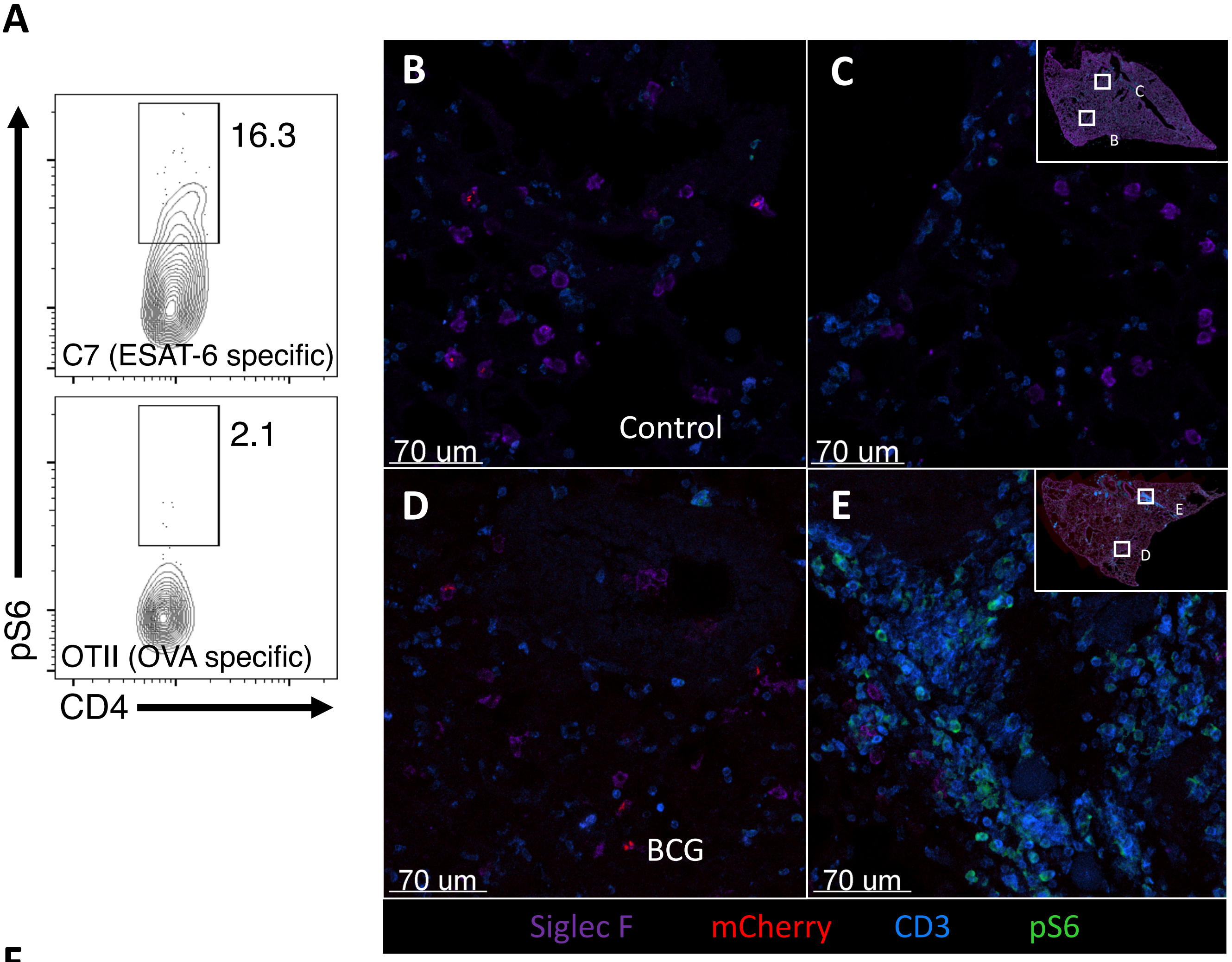


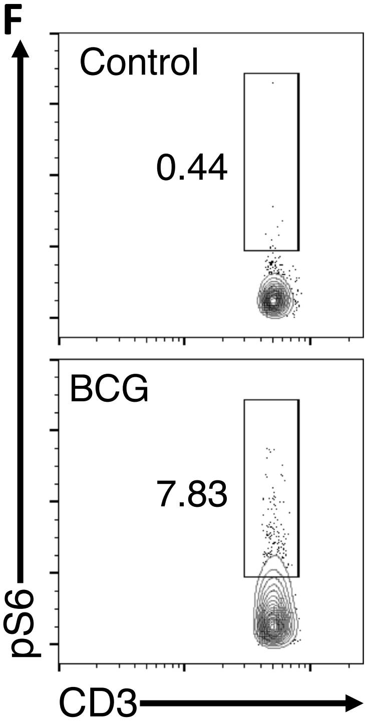


Supplemental Figure 2: Imaging and quantitative histocytometry of Mtb-infected lungs.
(A) Mtb-specific C7 and OVA-specific OTII CD4 T cells were stimulated *in vitro* to generate Th1 cells. These were then co-transferred into Mtb infected animals. Quantitative histocytometry was used to determine pS6+ C7 and OTII cells. Representative immunofluorescence images of control (B and C) and BCG-immunized (D and E) lungs at D10. Inset shows the areas of the lungs where mCherry$^{+}$ infected cells were found (B and D), as well as areas distal to the infection (C and E). (F) Representative flow plot showing the proportion of pS6$^{+}$ CD4 T cells from control and vaccinated lungs at D10.
